# Detection of Multiple Types of Cancer Driver Mutations Using Targeted RNA Sequencing in NSCLC

**DOI:** 10.1101/2021.08.25.457723

**Authors:** Sheng Ju, Zihan Cui, Yuayuan Hong, Xiaoqing Wang, Weina Mu, Zhuolin Xie, Xuexia Zeng, Lin Su, Qi Zhang, Xiaofeng Song, Songxia You, Ruixin Chen, Weizhi Chen, Xuchun, Jun Zhao

## Abstract

Currently, DNA and RNA are used separately to capture different types of gene mutations. DNA is commonly used for the detection of SNVs, indels and CNVs; RNA is used for analysis of gene fusion and gene expression. To perform both DNA sequencing (DNA-seq) and RNA-seq, material is divided into two copies, and two different procedures are required for sequencing. Due to overconsumption of samples and experimental process complexity, it is necessary to create an experimental method capable of analyzing SNVs, indels, fusions and expression.

We developed an RNA-based hybridization capture panel targeting actionable driver oncogenes in solid tumors and corresponding sample preparation and bioinformatics workflows. Analytical validation with an RNA standard reference containing 16 known fusion mutations and 6 SNV mutations demonstrated a detection specificity of 100.0% [95% CI 88.7%~100.0%] for SNVs and 100.0% [95% CI 95.4%~100.0%] for fusions. The targeted RNA panel achieved a 0.73-2.63 copies/ng RNA lower limit of detection (LOD) for SNVs and 0.21-6.48 copies/ng RNA for fusions. Gene expression analysis revealed a correlation greater than 0.9 across all 15 cancer-related genes between the RNA-seq results and targeted RNA panel.

Among 1253 NSCLC FFPE tumor samples, multiple mutation types were called from DNA- and RNA-seq data and compared between the two assays. The DNA panel detected 103 fusions and 21 METex14 skipping events; 124 fusions and 26 METex14 skipping events were detected by the target RNA panel; 21 fusions and 4 METex14 skipping events were only detected by the target RNA panel. Among the 173 NSCLC samples negative for targetable mutations by DNA-seq, 15 (15/173, 8.67%) showed targetable gene fusions that may change clinical decisions with RNA-seq. In total, 226 tier I and tier II missense variants for NSCLC were analyzed at genomic (DNA-seq) and transcriptomic (RNA-seq) levels. The positive percent agreement (PPA) was 97.8%, and the positive predictive value (PPV) was 98.6%. Interestingly, variant allele frequencies were generally higher at the RNA level than at the DNA level, suggesting relatively dominant expression of mutant alleles. PPA was 97.6% and PPV 99.38% for EGFR 19del and 20ins variants.

We also explored the relationship of RNA expression with gene copy number and protein expression. The RPKM of EGFR transcripts assessed by the RNA panel showed a linear relationship with copy number quantified by the DNA panel, with an R of 0.8 in 1253 samples. In contrast, MET gene expression is regulated in a more complex manner. In IHC analysis, all 3+ samples exhibited higher RPKM levels; IHC level of 2+ and below showed lower RNA expression.

Parallel DNA- and RNA-seq and systematic analysis demonstrated the accuracy and robustness of the RNA sequencing panel in identifying multiple types of variants for cancer therapy.

## INTRODUCTION

Kinase fusions, as drug targets, have been observed in various malignant tumors. Genes that have been reported to be involved in kinase fusion in non-small-cell lung cancer (NSCLC) include ALK, RET, ROS1, NTRK, NGR1, FGFR, EGFR, MET, etc. The development of targeted therapy has benefited lung cancer patients and improved their survival rate^1–3^. Examinations such as fluorescence in situ hybridization (FISH)/immunohistochemistry (IHC) and reverse transcription-polymerase chain reaction (RT-PCR) are commonly performed for the diagnosis of gene fusion. However, in addition to their low-throughput nature, FISH and IHC have other limitations, such as the lack of partner gene and fusion breakpoint information, and RT-PCR can only detect known fusions. As next-generation sequencing (NGS) technology has developed, it has been increasingly used in clinical diagnosis due to its high throughput nature and ability to identify breakpoints and fusion partners and discover new fusions^4^. NGS detection of fusion mutations includes two platforms: a DNA-based platform and an RNA-based platform. At the DNA level, fusion breakpoints usually occur in lengthy intron regions, and different patients may have different fusion breakpoints, so PCR-based detection is difficult to achieve. The DNA panel-based capture detection probe needs to cover exons and lengthy, multiplicity intron regions, and the probe design is technically difficult and expensive. With the RNA panel capture method, the probe only covers the exon region, and it is less technically difficult to detect fusion genes at the RNA level than at the DNA level. Jeong-Sun Seo et al found that fusion mutations activate genes with low expression under normal conditions and produce a large number of mRNAs^5^. The RNA platform can detect fusions more sensitively than the DNA platform. In addition, due to the existence of transcriptional regulation processes, RNA or protein detection is needed to determine whether DNA fusions are transcribed or translated. In summary, RNA platform-based detection of fusions has exceptional advantages.

RNA is transcribed from DNA and then translated to a protein. As an intermediate between DNA and protein, RNA is more reliable than DNA for predicting the function of proteins or interpreting the activity of cellular pathways. However, RNA is rarely used for detecting somatic mutations such as single-nucleotide variations (SNVs) or insertions/deletions (indels) because the accuracy of detection can be affected. For example, RNA testing may 1) have increased alignment errors near RNA splicing sites; 2) have an increased error rate in the process of reversed transcription; 3) fail to detect non-expressed or low-level variations due to different levels of expression; and 4) confuse RNA editing sites with mutations. It has been demonstrated that in tumor tissues RNA sequencing (RNA-seq) of SNVs in BRAF and RAS genes successfully detected 94% of the mutations with a frequency >10% detected by DNA exome sequencing; however, only 11% of the mutations with a variant allele frequency (VAF) of 5-10% were detected by RNA-seq^6^.

Currently, DNA is commonly used for SNV, indel and copy number variation (CNV) analysis, and RNA is used for fusion and gene expression analysis. Products on the market that claim to be able to use DNA and RNA to detect various types of mutations at the same time generally use capture or amplification methods. The capture method builds DNA and RNA libraries separately and captures DNA and RNA separately; in the amplification method, DNA and RNA are amplified separately, and then the amplified products are mixed together to build a library. In essence, these technical methods still require two types of experimental procedures, so the operation is complicated, time-consuming and cost-consuming. Therefore, it would be very meaningful to develop an experimental method suitable for SNV, indel, fusion, and expression analysis in parallel.

We reviewed the literature and used databases to identify studies related to gene fusion and the use of targeted drugs in solid tumors, and we selected 30 fusion genes and 6 housekeeping genes related to solid tumors, except for lymphoma, to design the panel.

This study used 1253 clinical NSCLC FFPE samples to perform DNA panel and RNA panel detection at the same time, with DNA panel detection results as a control, to explore the feasibility and accuracy of the RNA panel for SNV, indel, fusion and CNV detection. To date, this is the largest NSCLC tissue series for the detection of multiple types of cancer driver mutations by parallel DNA-seq and RNA-seq. The concordance of SNV, indel, fusion and CNV mutation data at the genomic DNA and transcriptomic RNA levels was systematically analyzed, and we preliminarily proved that the cost-controllable and reproducible RNA-targeted sequencing method can be applied for the precise detection of mutations related to targeted drugs for patients with NSCLC.

## MATERIALS AND METHODS

### MATERIALS

The samples used in this experiment were formalin-fixed, paraffin-embedded (FFPE) sections. All FFPE sections were sliced to 5-10-μm thick to assess the proportion of tumors and extract RNA.

### METHODS

#### RNA panel experiments

RNA was extracted from FFPE samples using the MagPure FFPE RNA/DNA Kit (Magen, China) with a 48 Automated Nucleic Acid Extractor (Magen, China) according to the manufacturer’s recommendations. The RNA concentration was measured by the Qubit RNA HS Assay Kit (Thermo Scientific, USA).

RNA (50-300 ng) from FFPE sections was used for library construction using mRNA-seq Lib Prep Module for Illumina (ABclonal, China). The RNA was processed through first-strand synthesis, second-strand synthesis, end repair, A-tailing, and adaptor (IDT, USA) ligation. Libraries were enriched with 15 cycles of PCR. The yield of libraries was measured with the Equalbit dsDNA HS Assay Kit (Vazyme, China). The size of libraries was assessed with the HT DNA NGS 3K Reagent Kit (Bluescape, China).

The RNA panel was designed and synthesized by IDT. Hybridization and washing were conducted according to the manufacturer’s protocol of the xGen Hybridization and Wash Kit and xGen Universal Blockers-TS Mix-96rxn (IDT, USA). The capture libraries were enriched with PCR. The captured products were sequenced for 150 bp paired-end sequencing and run on an Illumina NovaSeq 6000 instrument according to the manufacturer’s recommendations.

#### RNA panel data analysis

Trimmomatic-0.36 was used to remove low-quality reads and adaptor sequences, the high-quality sequence reads were retained, STAR was used to compare sequence reads with the reference genome, and sequence quality evaluation was performed. Only those reads that pass the quality requirement can be used for downstream analysis. Downstream analysis included gene expression analysis, gene fusion analysis, fusion expression analysis and SNV and indel analysis.

##### 1) Gene expression analysis

According to the sequence alignment results and the annotation files of the reference genome, the reads per kilobase per million mapped reads (RPKM) method was used to evaluate gene expression.

##### 2) Gene fusion analysis

FusionMap was used to identify gene fusions. The first step is to initially obtain the results of gene fusion. The second step was to filter the fusion results. The filter included fusion genes of the same gene family, fusion genes of the same paralogous group (defined by Ensembl v74), and fusion genes derived from the same gene model. The third step was to filter the results according to the fusion threshold to obtain the final result.

##### 3) Fusion expression analysis

The fusion expression level was obtained by using the housekeeping gene to correct the expression level of the fusion gene. The calculation formula is as follows:

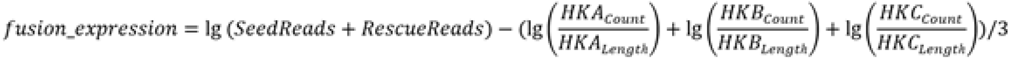

##### 4) SNV and indel analysis

The sequencing data were aligned to the reference genome (hg19), and a bam file was obtained. VarDict caller was used to capture the mutation sites and indels, and the result file was in VCF format. The VCF files were annotated by ANNOVAR, and Transvar annotations were used for some inaccurate sites. Plus and minus strands were corrected, and the number of reads and frequency of variants were counted. The internal evidence database was used to filter annotations and support transcript selection. The SNV and indel threshold were used to filter and obtain the final result.

#### RNA-seq

RNA was assessed using the Agilent 2100 Bioanalyzer system assay. Then, 50 ng of total RNA was used with the SMARTer Stranded Total RNA-Seq Kit v2 (Takara) in accordance with the low-throughput protocol. This kit incorporates SMART^®^ (switching mechanism at the 5′ end of RNA template) cDNA synthesis technology and generates Illumina-compatible libraries via PCR amplification, avoiding the need for adaptor ligation. After PCR enrichment and purification of adaptor-ligated fragments, the library with adaptors was determined with the Qubit dsDNA HS (High Sensitivity) Assay Kit (Thermo Fisher, USA) and assessed using the Agilent 2100 Bioanalyzer system assay. Then, RNA-seq was performed using the Illumina NovaSeq 6000 Sequencing System.

To ensure data quality, raw reads were preprocessed by removing low-quality sequences, de-junction contamination, rRNA signals, etc. to obtain high-quality sequences (clean reads), and all subsequent analyses were based on clean reads. Reference gene and genome annotation files were downloaded from the ENSEMBL website (http://www.ensembl.org/index.html). Clean data were aligned to the reference genome by HISAT2 (http://ccb.jhu.edu/software/hisat2/index.shtml). HTSeq (http://www.huber.embl.de/users/anders/HTSeq/doc/overview.html) was used to estimate the expression level of each gene. The quantification of gene expression was performed using the RPKM method.

#### DNA panel capture

Genomic DNA was extracted from FFPE samples using the MagPure FFPE DNA Kit B (Magen, China). Peripheral blood lymphocyte (PBL) DNA was extracted by the TGuide S32 Magnetic Blood Genomic DNA Kit (Tiangen, China). The DNA concentration was measured by the Qubit dsDNA HS (High Sensitivity) Assay Kit (Thermo Fisher, USA). The quality of DNA was assessed by an Agilent 2100 BioAnalyzer (Agilent, USA).

DNA (50-300 ng) from FFPE samples was sheared with Covaris LE220 according to the recommended settings for ~200 bp fragments, and the fragmented DNA was input for library construction by the KAPA Hyper Preparation Kit (Kapa Biosystems, USA) according to the manufacturer’s instructions. All libraries were quantified by the AccuGreen High Sensitivity dsDNA Quantitation Kit (Biotium, USA), and the size of the libraries was determined on the Agilent Bioanalyzer 2100 (Agilent, USA).

Targeted region selection was performed with the xGen Hybridization and Wash Kit (IDT). The panel was designed of 543 genes frequently mutated in common solid tumors. The captured product was sequenced using 150-bp paired-end runs on an Illumina NovaSeq 6000 instrument according to the manufacturer’s recommendations.

The generated sequencing reads were treated with Trimmomatic for adaptor trimming and filtering out low-quality bases. The obtained reads were aligned to the hg19 human genome reference using BWA aligner v 0.7.12. The aligned and sorted BAM files were then removed from duplicated reads by using Picard 2.2.1. GATK v 3.8 was used to perform local realignment around potential small indels and base recalibration for next step mutation calling procedures. We used Verdict v1.5.1 to detect SNV and small indels and freebayes v1.1.0-44 to detect complex variants. Tumor-normal paired sample calling was performed during the mutation calling procedure to filter out personal germline mutations.

DNA copy number alterations were detected by using CNV kit v.0.8.5, which compares tumor samples with a pool of all selected CNV-negative patient FFPE samples. This algorithm considers both on- and off-target sequencing reads and applies a series of corrections for extraneous variability sources, such as GC content, target footprint size and spacing, to improve accuracy in copy number calling.

#### IHC

Following the standard IHC protocol, a single primary antibody reagent was used for detection with the SP method (Zhongshan Jinqiao PV8000 Two-step Universal Kit); the paraffin-embedded tissue sections were deparaffinized with xylene, and the tissue sections were washed with PBS and dehydrated with graded alcohols. Then, pH 8.0 EDTA was applied, and the sample was autoclaved for antigen retrieval; endogenous peroxidase activity was quenched with 3% H_2_O_2_. The sample was washed with PBS and then distilled water. The slide was then incubated with the primary antibody. Then, the slide was incubated with the secondary antibody. After washing, the slides were incubated with 3,3′-diaminobenzidine tetrahydrochloride (DAB) and washed in tap water immediately after color development. Then, the slides were counterstained with hematoxylin and differentiated with hydrochloric acid. The repair solution was applied for counterstaining (blue) and dehydrated with graded alcohols. The slides were mounted with transparent xylene and neutral gum and then observed under an optical microscope.

## RESULTS

### Introduction of RNA panel

We reviewed the literature and used databases to identify studies that were related to gene fusion and the use of targeted drugs in solid tumors, and we selected 30 fusion genes and 6 housekeeping genes related to solid tumors, except for lymphoma, to design the panel. The selected fusion genes included 18 genes that were related to targeted drugs, including ALK, BRAF, EGFR, ESR1, EWSR1, FGFR1, FGFR2, FGFR3, FGFR4, MET, NRG1, NTRK1, NTRK2, NTRK3, PDGFRA, PDGFRB, RET, and ROS1, as well as 12 partner genes. The 6 housekeeping genes that were selected were ciliary neurotrophic factor (CNTF), epilepsy progressive myoclonus type 2A (EPM2A), HEATR4, nucleolar protein 10 (NOL10), retinitis pigmentosa GTPase regulator interacting protein 1 (RPGRIP1), and SFT2 domain containing 3 (SFT2D3). The selected regions for the panel design were submitted to Integrated DNA Technologies (IDT). The completed panel, FUCAST, was 126 kb and contained 2133 probes.

RNA was extracted, and target sequencing was performed using the RNA panel. Data analysis showed that the map ratio was 0.966±0.069, and the target ratio was 0.715±0.121.

### Evaluation of fusion gene detection by RNA

RNA was extracted from SeraCare Formalin-Fixed, Paraffin-Embedded (FFPE) Tumor Fusion RNA Reference Material (tumor v2) and diluted 1:10, 1:20, 1:50, 1:75, 1:100 and 1:125 with GM12878 FFPE RNA. According to the tumor v2 fusion copy number supported by the manufacturer, the number of fusion transcript copies contained in dilutions can be calculated. Fifty nanograms of RNA from different serial dilution samples was used for library construction and RNA panel capture, and each dilution was repeated nine times. When the fusion mutant was detected in all nine repeats, the limit of detection (LOD) was defined. GM12878 FFPE RNA was used as a negative control to investigate the RNA fusion specificity, and no fusion was detected in the negative control.

To analyze the repeatability of the fusion detection by the RNA panel, the SeraCare FFPE Tumor Fusion RNA Reference Material (tumor v2) sample was diluted 20X with GM12878 FFPE RNA. Fifty nanograms of diluted FFPE RNA was used for library construction and RNA panel capture. Under the same experimental conditions, each sample was repeatedly tested 3 times. The analysis of the 16 fusion types showed that the repeatability of the RNA fusion detection reached 100% (RNA = 1.03-3.24 copies/ng).

### Validation of fusion gene detection in clinical samples

#### Patient cohort and clinical data

To validate the reliability of the RNA panel to detect gene mutations, next-generation sequencing (NGS) was performed on a large number of tissue samples (n = 1270) at both the DNA level and the RNA level. The samples were collected from patients with non-small-cell lung cancer (NSCLC) between May 15, 2020, and March 08, 2021. According to RNA panel quality control standards (target reads>=2 M bp, intron + intergenic rate <30%, average unique counts of 3 housekeeping genes (EPM2A, RPGRIP1 and SFT2D3) > 100), we excluded 17 samples and kept only 1253 samples for further testing.

Among the NSCLC cases involved in the study, lung adenocarcinoma accounted for 78.45% of cases (n =983), squamous cell carcinoma (SCC) accounted for7.66% of cases (n = 96) and other types of lung cancer accounted for 13.89% of cases (n =174). In terms of smoking history, smokers accounted for 37.03% of patients (n =464), and never-smokers accounted for 59.46% of patients (n = 745). Both DNA-based testing and RNA-based testing showed that nonsmokers were more likely to harbor gene fusions than smokers (DNA: 80.58% vs 12.62%; RNA: 80.65 vs 13.71%) (see Table 2).

#### Detection of drug-targetable mutations

Gene mutations that are highly associated with NSCLC treatment can be categorized into three types: SNVs (involving ALK, BRAF, EGFR, ERBB2, KRAS, MET, ROS1), copy number variation (involving MET and ERBB2), and gene fusion (involving ALK, RET, ROS1). In the study, among 1253 patients with NSCLC, 1080 patients (86.19%) showed positive results for driver genes through the DNA panel, where the EGFR (59.35%) and KRAS (12.22%) genes were the driver genes with mutated at the highest frequencies (Fig. 1).

**Fig. 1.**
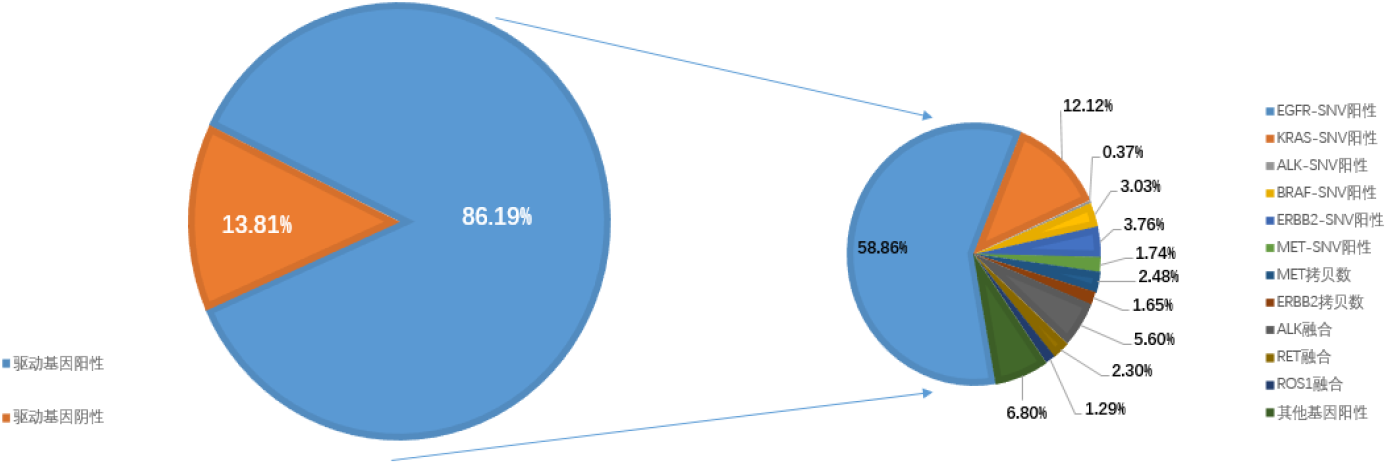
The proportion of each driver gene in the positive sample of driver genes

Moreover, 173 patients (173/1253, 13.80%) with NSCLC were identified as driver-negative by DNA sequencing (DNA-seq). However, through further RNA sequencing (RNA-seq), 15 of them were found to be fusion-positive, which can predict the efficacy of potential targeted drugs. The gene fusions that were detected involved ALK (n = 7), RET (n =3), METex14 skipping (n =3), ROS1 (n=1) and MET (n=1). The results demonstrated that compared to the DNA panel, the RNA panel can detect gene fusion more sensitively and can increase the possibility that patients are administered the appropriate targeted drugs.

#### The RNA panel identified more gene fusions than the DNA panel

One hundred and twenty-four cases were detected as fusion-positive in the RNA panel, whereas the DNA panel detected 103 of them. Through the RNA panel, we found ALK (68/1253, 5.43%), RET (29/1253, 2.31%), ROS1 (20/1253, 1.60%), MET (3/1253, 0.24%), FGFR2/3 (2/1253, 0.16%), and 2 cases with an uncommon fusion in NSCLC (BRAF, ESR1). For the DNA panel, 4.79% had ALK fusions (n = 60), 2.00% had RET fusions (n = 25), 1.36% had ROS1 fusions (n=17), and 0.08% had MET fusions (n = 1) (see Table 1 in the online Data Supplement). In this study, we noticed that the RNA panel was capable of identifying more patients who harbored gene fusion than the DNA panel (8 cases of ALK, 3 of ROS1, 4 of RET, 2 of MET, 2 of FGFR, and 1 of ESR1, BRAF); 21 cases of RNA fusions were not detected by the DNA panel (see Table 1 in the online Data Supplement). Possible reasons for the DNA panel not detecting fusions are the range of the panels. Two ROS1 fusions in exon 32 were detected by the RNA panel; the breakpoint position was expected to be located in intron 31, and a previous study demonstrated a large number of repeat components in intron 31. Hence, it is challenging to design a targeted panel for DNA^7^. As previously reported by Ying^8^, we also detected multiple genes or Intergenic involved in rearrangement at the DNA level, while fusion-positive patients were detected at the RNA level (see Fig. 1 in the online Data Supplement). RNA panel capture detection of fusions has advantages over DNA panels.

**Table 1.** The LOD of fusion detected by RNA panel

| Gene1 | Exon1 | Gene2 | Exon2 | LOD (copies/ng) |
| --- | --- | --- | --- | --- |
| KIF5B | Exon24 | RET | Exon11 | 0.21 |
| NCOA4 | Exon8 | RET | Exon12 | 1.80 |
| ETV6 | Exon5 | NTRK3 | Exon15 | 1.60 |
| TPM3 | Exon8 | NTRK1 | Exon9 | 0.32 |
| LMNA | Exon2 | NTRK1 | Exon10 | 1.21 |
| SLC45A3 | Exon1 | BRAF | Exon8 | 0.33 |
| TMPRSS2 | Exon1 | ERG | Exon2 | 1.03 |
| EML4 | Exon13 | ALK | Exon20 | 2.30 |
| PAX8 | Exon9 | PPARG | Exon2 | 6.48 |
| FGFR3 | Exon17 | TACC3 | Exon11 | 0.49 |
| FGFR3 | Exon17 | BAIAP2L1 | Exon2 | 0.49 |
| CD74 | Exon6 | ROS1 | Exon34 | 3.82 |
| EGFR | Exon24 | SEPT14 | Exon10 | 1.66 |
| EGFR | Exon1 | EGFR | Exon8 | 0.31 |
| MET | Exon13 | MET | Exon15 | 0.83 |

**Table 2.**
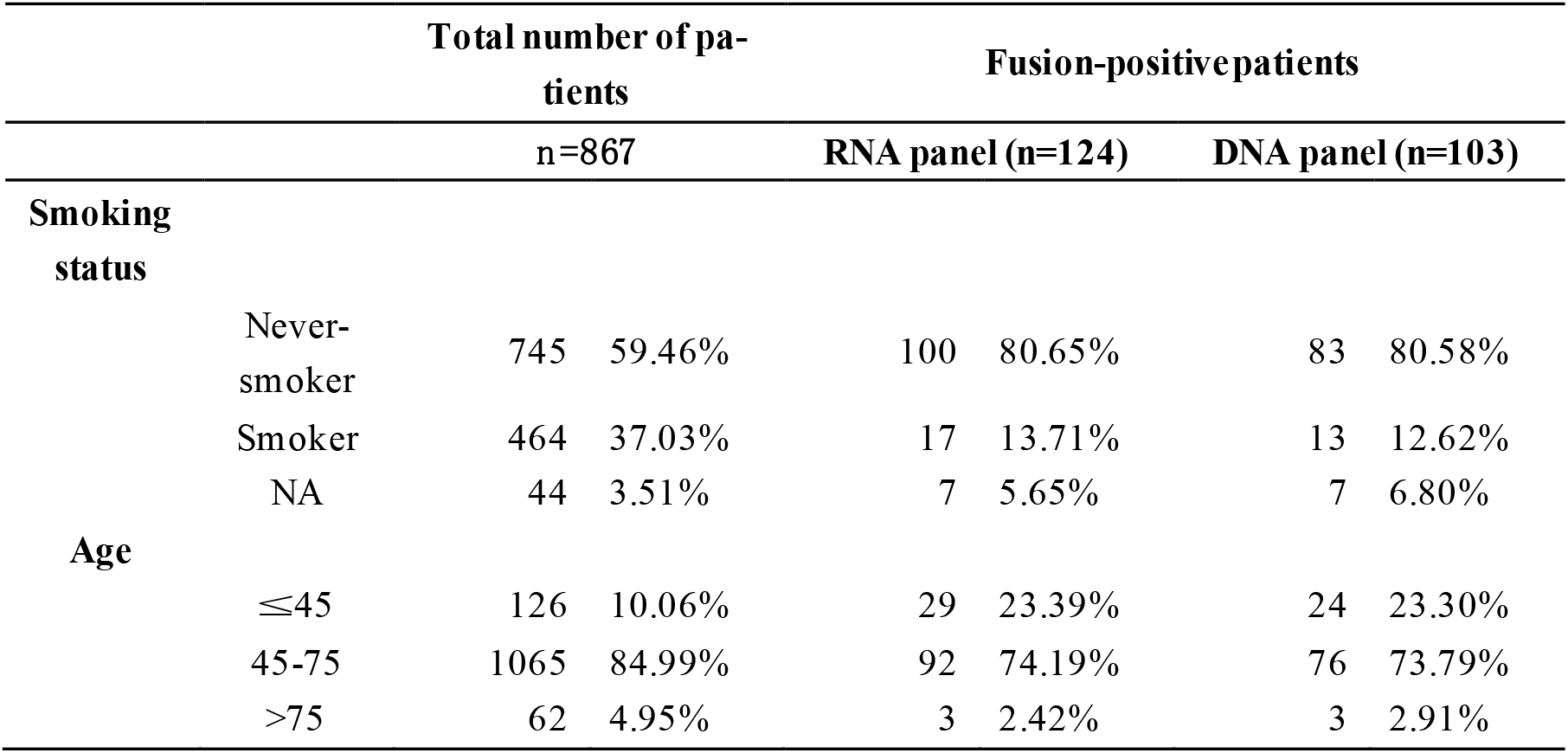
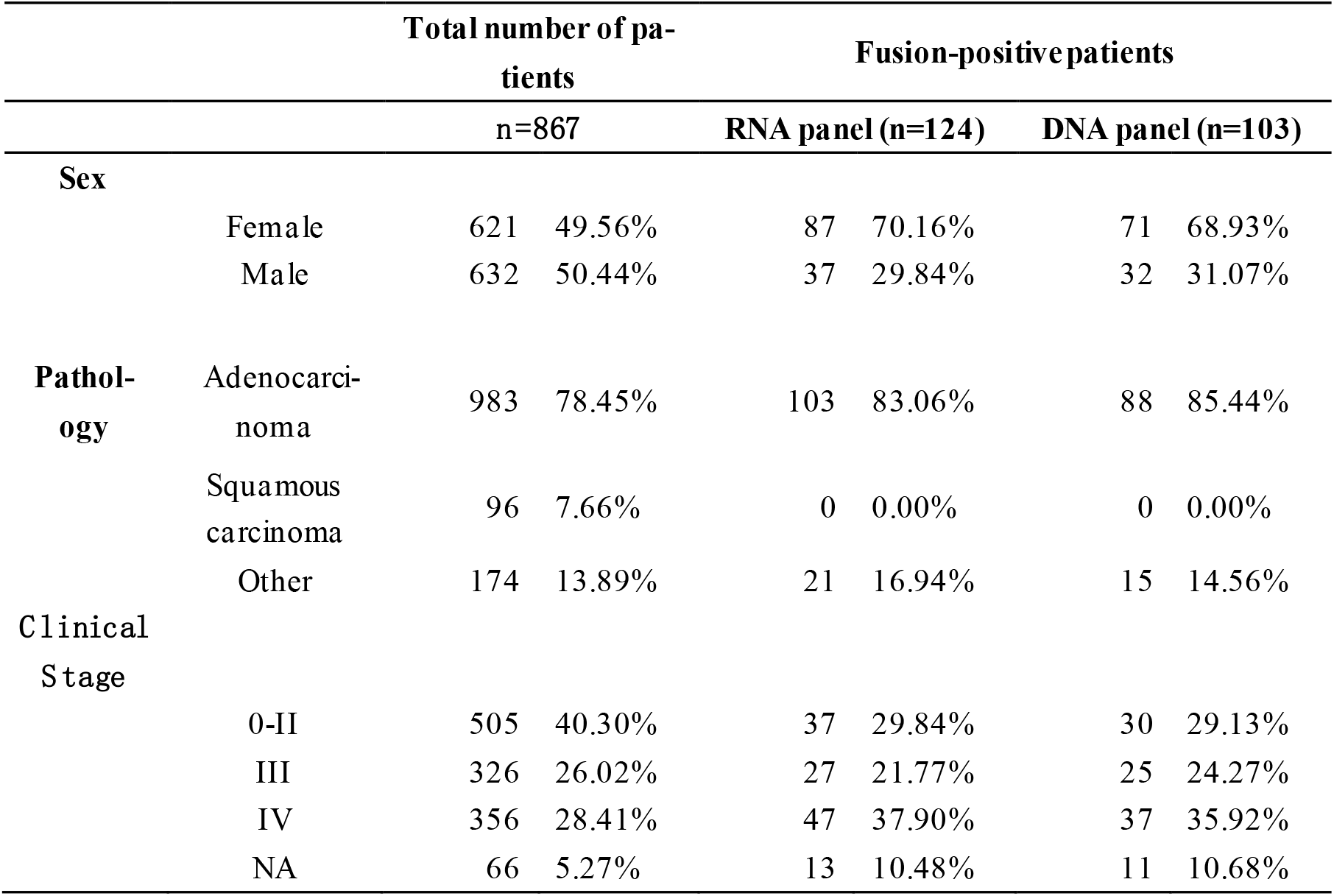
Clinical information

#### ALK fusion expression analysis

There are more than 20 subtypes of EML4-ALK fusion variants, and the most common are V1 and V3a/b. The RNA panel detected 23 and 28 cases, respectively, accounting for 79.68% (51/64) of all ELM4-ALK fusions; statistical RNA panel. The relative expression levels of the two types of fusion subtypes were tested (see methods), and it was found that when the fusion mutation was V1 type (n=28), the fusion gene expression level was significantly higher than that of V3a /b (n=13 and n=15, respectively; P<0.001). For verification, if there is enough remaining template, we use droplet digital PCR (ddPCR) to verify the expression level of the fusion and found the same trend^9^, as shown in Fig. 2. (see Table S2 in the online Data Supplement)

**Fig. 2.**
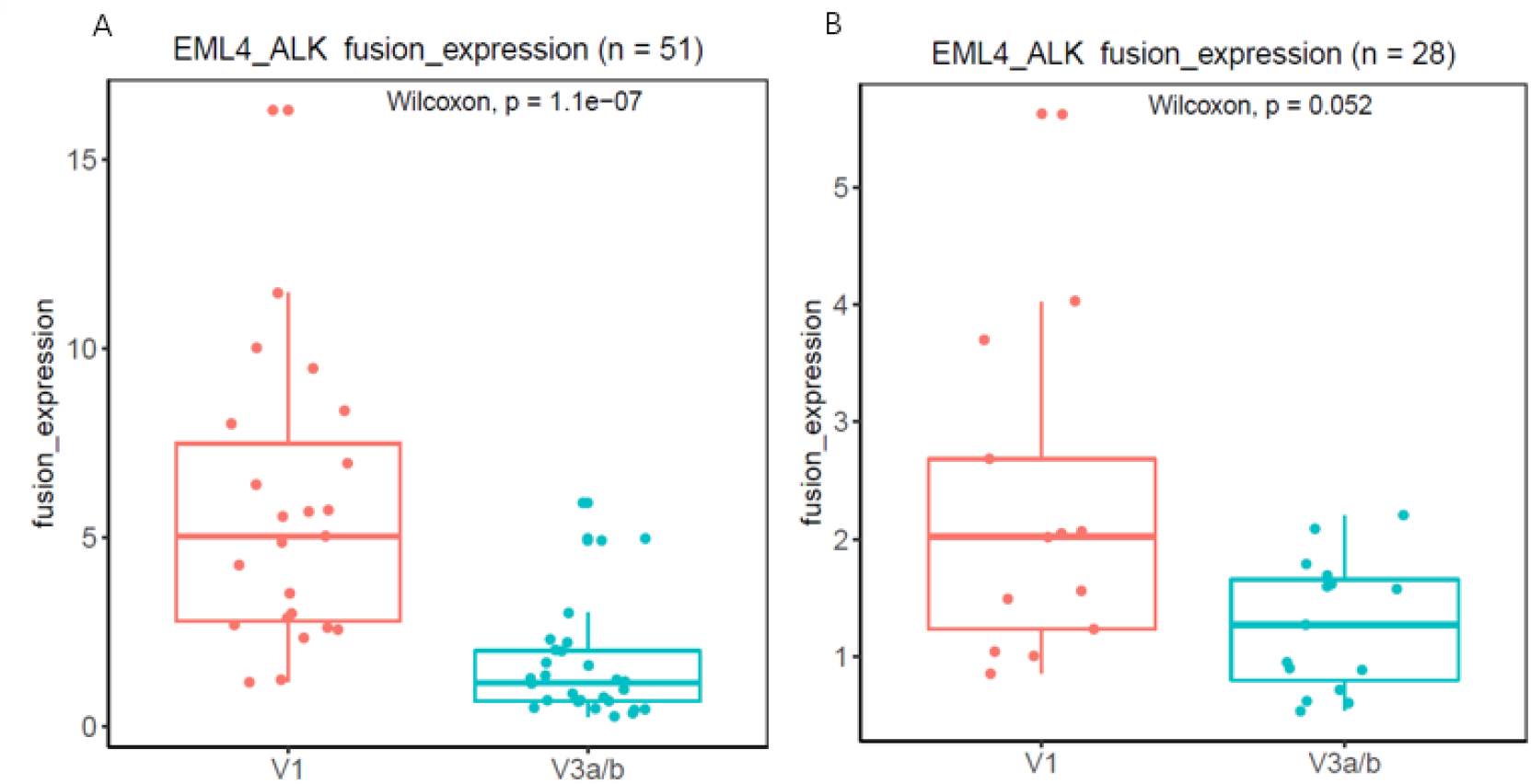
Fusion gene expression level (corrected by the proportion of tumor cells judged by standard pathology). (A) RNA panel detection of the fusion gene expression level. (B) ddPCR detection of the relative expression level of the fusion gene.

#### METex14 skipping

Among the 26 patients who had METex14 skipping mutations detected through the RNA panel, 6 patients (23.08%, 6/26) also had classic splicing mutations detected by the DNA panel, 16 patients (56.25%, 9/16) had noncanonical splicing mutations at ±20 bp splicing loci, and 6 had no splicing mutations. Further analysis of the original data revealed that 1 patient harbored the MET c.3028+5A>T mutation at the splicing region. However, the single-nucleotide variation (SNV) was distant from the splicing locus. No related mutations at the splicing region were detected in the other 5 patients (see Table S3 in the online Data Supplement)

#### Evaluation of SNV detection by RNA

A total of 226 loci that were either sensitive or resistant to targeted drugs overlapped in both the RNA and DNA panels. The loci were located in 11 genes, including ALK, BRAF, EGFR, ESR1, FGFR2, FGFR3, MET, NTRK1, NTRK3, PDGFRA and ROS1. Six types of SNV mutations were randomly selected (Table 3) and synthesized by in vitro transcription. They were mixed together as a standard pool, and the copy number of each mutation was quantified by ddPCR. The standard pool was serially diluted with GM12878 RNA, and four different concentration pools were obtained. Fifty nanograms of RNA from different serial dilution samples was used for library construction and RNA panel capture, and each dilution was repeated three times. When the SNV mutant was detected in all three repeats, the LOD was defined. GM12878 RNA was used as a negative control to investigate RNA SNV specificity.

**Table 3.** Six RNA SNV mutants synthesized by in vitro transcription

| Gene | Chr | Transcript | Exon | Amino acid variant | CDS mutation |
| --- | --- | --- | --- | --- | --- |
| FGFR2 | Chr10 - | NM_000141 | 7 | p.P253R | c.758C>G |
| FGFR3 | Chr4 + | NM_000142 | 7 | p.R248C | c.742C>T |
| NTRK3 | Chr15 - | NM_001012338 | 16 | p.G623R | c.1867G>A |
| ALK | Chr2 - | NM_004304 | 23 | p.L1196M | c.3586C>A |
| ALK | Chr2 - | NM_004304 | 23 | p.G1202R | c.3604G>A |
| ROS1 | Chr6 - | NM_002944 | 38 | p.D2033N | c.6097G>A |

The SNV mutant reads for the 226 loci in the negative control GM12878 cell line RNA analysis was ≤1. Based on the following formula: threshold = average value ± 3 standard deviations, the RNA SNV read threshold was 2. Based on the threshold, the LOD of the 6 SNV mutations was defined (Table 4).

**Table 4.** The LOD of the six RNA SNV mutants

| SNV | LOD (copies/ng) |
| --- | --- |
| FGFR2 c.758C>G | 1.41 |
| FGFR3 c.742C>T | 2.63 |
| NTRK3 c.1867G>A | 1.57 |
| ALK c.3586C>A;c.3604G>A | 0.99 |
| ROS1 c.6097G>A | 0.73 |

#### Evaluation of SNV detection by the RNA panel in clinical samples

The results of the parallel DNA panel and RNA panel showed that in the 1253 NSCLC samples, the correlation coefficient of the SNV variant frequency detected by the DNA and RNA was 0.80 (R= 0.80). The SNV variant frequency detected by the RNA panel was higher than that detected by the DNA panel (Fig. 3). This finding was confirmed by ddPCR (Table 5) and was consistent with that of a previous study^6^. There were two mutants whose DNA variant frequency was much higher than that observed with the RNA panel. They were located at the last base of METex14, existed at the DNA level but were lost at the RNA level due to MET ex14 skipping.

**Fig. 3.**
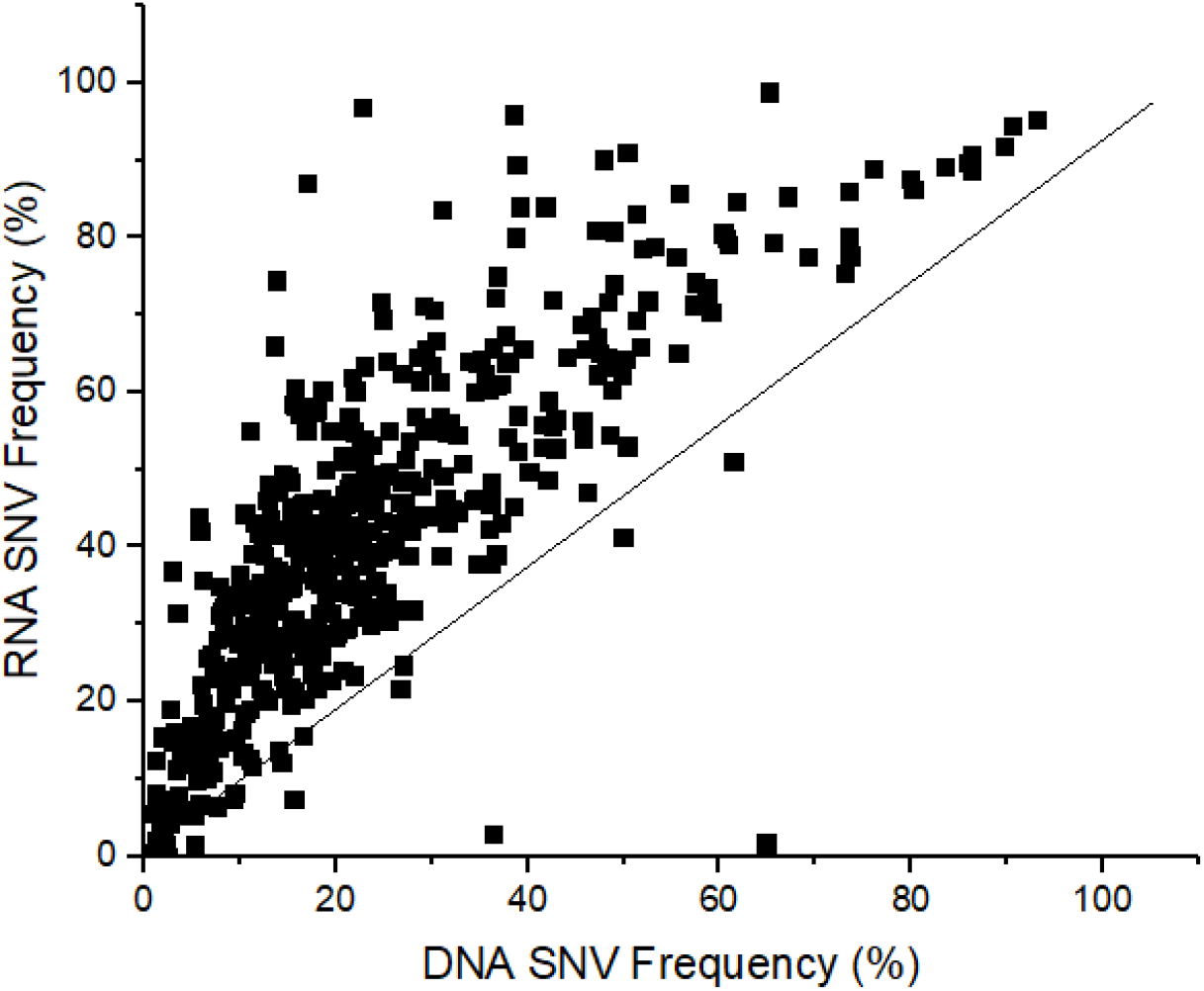
The correlation between the DNA SNV frequency and RNA SNV frequency

**Table 5.** SNVs frequency confirmed by ddPCR

| Sample ID | Gene | Mutation_p | DNA panel freq (%) | RNA panel freq (%) | DNA ddPCR freq (%) | RNA ddPCR freq (%) |
| --- | --- | --- | --- | --- | --- | --- |
| P210904 | EGFR | p.L858R | 3.22 | 16.18 | 3.50 | 16.40 |
| P210376 | EGFR | p.T790M | 1.81 | 15.47 | 1.72 | 14.50 |
| P210911 | EGFR | p.L858R | 2.67 | 19.00 | 2.13 | 18.50 |
| P222742 | EGFR | p.L858R | 5.79 | 42.03 | 6.30 | 42.00 |
| P223889 | EGFR | p.E709A | 6.04 | 35.77 | 4.73 | 28.30 |

The 1253 NSCLC samples were divided into two groups: 392 were used as the threshold training set, and 861 were used as the threshold verification set. According to the performance of the 392 samples under different threshold conditions of RNA SNV, the best threshold standard is variant allele frequency (VAF)>= 1.5% and mutation support reads>=10. Under this threshold, the RNA SNV detection performance of clinical samples in the training set was positive percent agreement (PPA)=97.87%, negative percent agreement (NPA)=100.00%, and positive predictive value (PPV)=99.28% (Table 6). The clinical sample threshold was higher than the threshold from the GM12878 cell line RNA analysis (mutation support reads>=2). When using the threshold from the GM12878 RNA analysis, the clinical samples showed more false positive results, indicating that the cell line negative control cannot reflect the situation of the clinical samples.

**Table 6.** The performance of RNA SNVs under different thresholds in the training set

| Mutant Frequency | Support Reads | TP | FN | TN | FP | PPA | PPV |
| --- | --- | --- | --- | --- | --- | --- | --- |
| 0.00% | 2 | 139 | 2 | 87506 | 945 | 98.58% | 12.82% |
| 0.00% | 5 | 139 | 2 | 87842 | 609 | 98.58% | 18.58% |
| 0.00% | 10 | 139 | 2 | 88222 | 229 | 98.58% | 37.77% |
| 0.00% | 15 | 138 | 3 | 88361 | 90 | 97.87% | 60.53% |
| 1.00% | 2 | 138 | 3 | 88405 | 46 | 97.87% | 75.00% |
| 1.00% | 5 | 138 | 3 | 88436 | 15 | 97.87% | 90.20% |
| 1.00% | 10 | 138 | 3 | 88449 | 2 | 97.87% | 98.57% |
| 1.00% | 15 | 138 | 3 | 88449 | 2 | 97.87% | 98.57% |

Table 6. Continued
| Mutant Fre-<br>quency | Support<br>Reads | TP | FN | TN | FP | PPA | PPV |
| --- | --- | --- | --- | --- | --- | --- | --- |
| 1.50% | 2 | 138 | 3 | 88425 | 26 | 97.87% | 84.15% |
| 1.50% | 5 | 138 | 3 | 88438 | 13 | 97.87% | 91.39% |
| 1.50% | 10 | 138 | 3 | 88450 | 1 | 97.87% | 99.28% |
| 1.50% | 15 | 138 | 3 | 88450 | 1 | 97.87% | 99.28% |

In the 861 validation set samples, according to the clinical sample threshold standard, the performance results obtained were PPA=97.88%, NPA=100.00%, and PPV=98.66%. A total of 363 of the 371 positive mutations detected by DNA could be detected by RNA. Analysis of the 8 missed mutations (Table 7), of which 1 case was a mutation with a DNA mutation frequency of >5%, and the remaining 7 mutations had a mutation frequency of <3%. Among them, 4 cases had mutant molecules at the RNA level, but they were missed because of the RNA SNV threshold. The distribution of positive mutations in DNA was EGFR (91.68%), BRAF (5.97%), ALK (0.72%), MET (0.90%) and FGFR3 (0.36%). The mutations of EGFR gene are mainly L858R (60.75%). 5 mutations were only detected in RNA (Table 8). P211694 was randomly selected for ddPCR verification, and the ddPCR results were consistent with the NGS results.

**Table 7.**
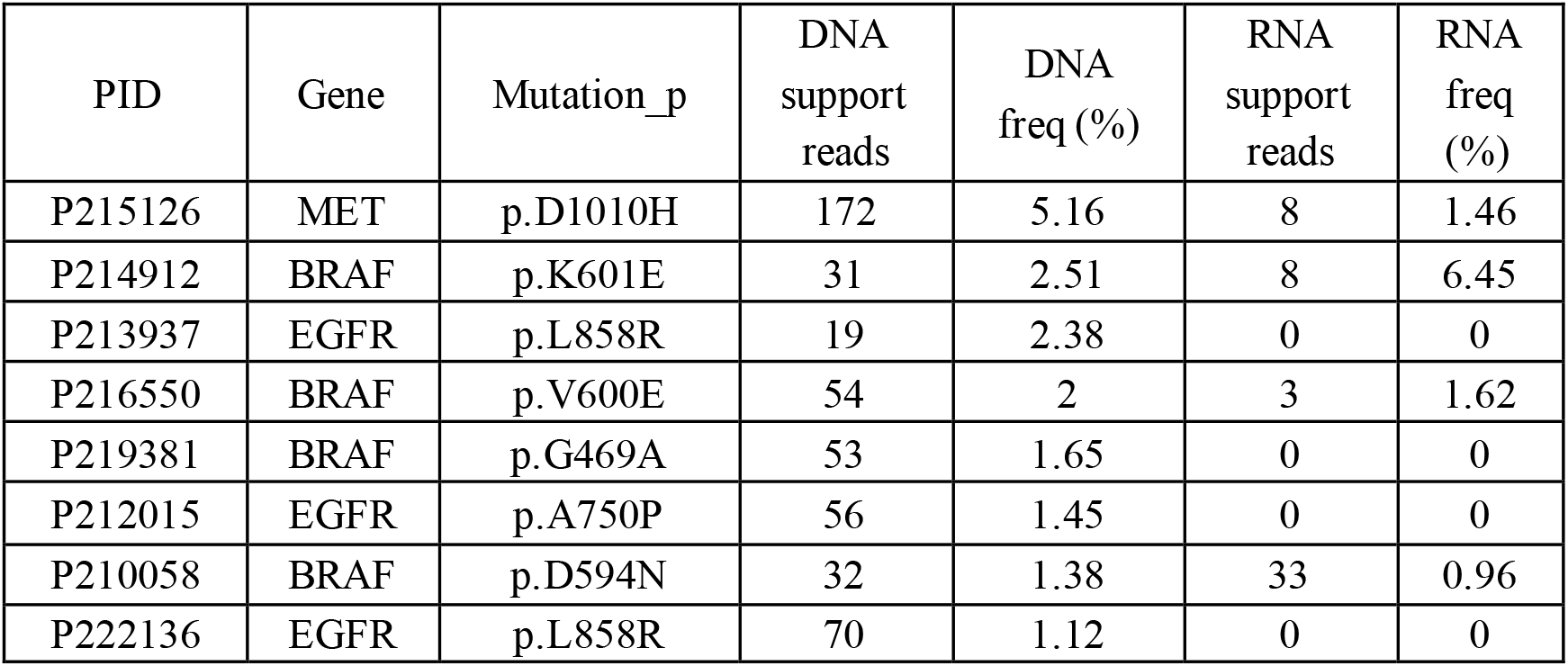
RNA missed SNV

| PID | Gene | Mutation_p | DNA<br>support<br>reads | DNA<br>freq (%) | RNA<br>support<br>reads | RNA<br>freq<br>(%) |
| --- | --- | --- | --- | --- | --- | --- |
| P215126 | MET | p.D1010H | 172 | 5.16 | 8 | 1.46 |
| P214912 | BRAF | p.K601E | 31 | 2.51 | 8 | 6.45 |
| P213937 | EGFR | p.L858R | 19 | 2.38 | 0 | 0 |
| P216550 | BRAF | p.V600E | 54 | 2 | 3 | 1.62 |
| P219381 | BRAF | p.G469A | 53 | 1.65 | 0 | 0 |
| P212015 | EGFR | p.A750P | 56 | 1.45 | 0 | 0 |
| P210058 | BRAF | p.D594N | 32 | 1.38 | 33 | 0.96 |
| P222136 | EGFR | p.L858R | 70 | 1.12 | 0 | 0 |

**Table 8.** SNVs detected only in RNA

| Sample ID | Gene | Mutation_p | DNA support reads | DNA freq (%) | RNA support reads | RNA freq (%) | DNA ddPCR freq (%) | RNA ddPCR freq (%) |
| --- | --- | --- | --- | --- | --- | --- | --- | --- |
| P222660 | ESR1 | p.D538G | 0 | 0 | 19 | 11.59 | NA | NA |
| P218528 | EGFR | p.L858R | 0 | 0 | 16 | 6.93 | NA | NA |
| P211694 | EGFR | p.G598V | 0 | 0 | 19 | 6.55 | 0 | 5.4 |
| P218876 | EGFR | p.L858R | 0 | 0 | 23 | 2.31 | NA | NA |
| P210229 | EGFR | p.T790M | 33 | 0.75 | 121 | 1.96 | NA | NA |

#### Evaluation of insertion/deletion (indel) detection by the RNA panel in clinical samples

Indels that were either sensitive or resistant to targeted drugs overlapped in both the RNA and DNA panels. The loci were located in 11 genes, including ALK, BRAF, EGFR, PDGFRA and MET. The 1253 NSCLC samples were divided into two parts: 392 cases were used as the threshold training set, and 861 cases were used as the threshold verification set. According to the performance of RNA detection for the 392 samples under different threshold conditions, the best threshold was obtained: VAF>= 2% and mutation support reads>=2. Under this threshold, the RNA indel detection performance of clinical samples in the training set was PPA=100.00%, and PPV=100.00% (Table 9).

**Table 9.** Performance of RNA indel detection under different thresholds in the training set

| Mutant Frequency | Support Reads | TP | FN | FP | PPA | PPV |
| --- | --- | --- | --- | --- | --- | --- |
| 1.0% | 2 | 106 | 0 | 1 | 100.00% | 99.07% |
| 1.0% | 5 | 106 | 0 | 1 | 100.00% | 99.07% |
| 1.0% | 10 | 106 | 0 | 1 | 100.00% | 99.07% |
| 1.0% | 15 | 106 | 0 | 1 | 100.00% | 99.07% |
| 1.5% | 2 | 106 | 0 | 1 | 100.00% | 99.07% |
| 1.5% | 5 | 106 | 0 | 1 | 100.00% | 99.07% |
| 1.5% | 10 | 106 | 0 | 1 | 100.00% | 99.07% |
| 1.5% | 15 | 106 | 0 | 1 | 100.00% | 99.07% |
| 2.0% | 2 | 106 | 0 | 0 | 100.00% | 100.00% |
| 2.0% | 5 | 106 | 0 | 0 | 100.00% | 100.00% |
| 2.0% | 10 | 106 | 0 | 0 | 100.00% | 100.00% |
| 2.0% | 15 | 106 | 0 | 0 | 100.00% | 100.00% |

In the 861 validation set samples, according to the clinical sample threshold, the performance results obtained were PPA=97.56%, and PPV=99.38%. A total of 160 of the 164 positive mutations detected by DNA could be detected by RNA. There were four missed mutations (Table 10), and one of them had a DNA mutation frequency >5%; mutations were present at the RNA level but were missed because of the RNA indel threshold. The remaining three missed mutations were low-frequency mutations with a mutation frequency of <5%. The MET alteration was METex14 skipping, so there was no mutation at the RNA level. The RNA analysis detected one more mutation than the DNA analysis (Table 11).

**Table 10.**
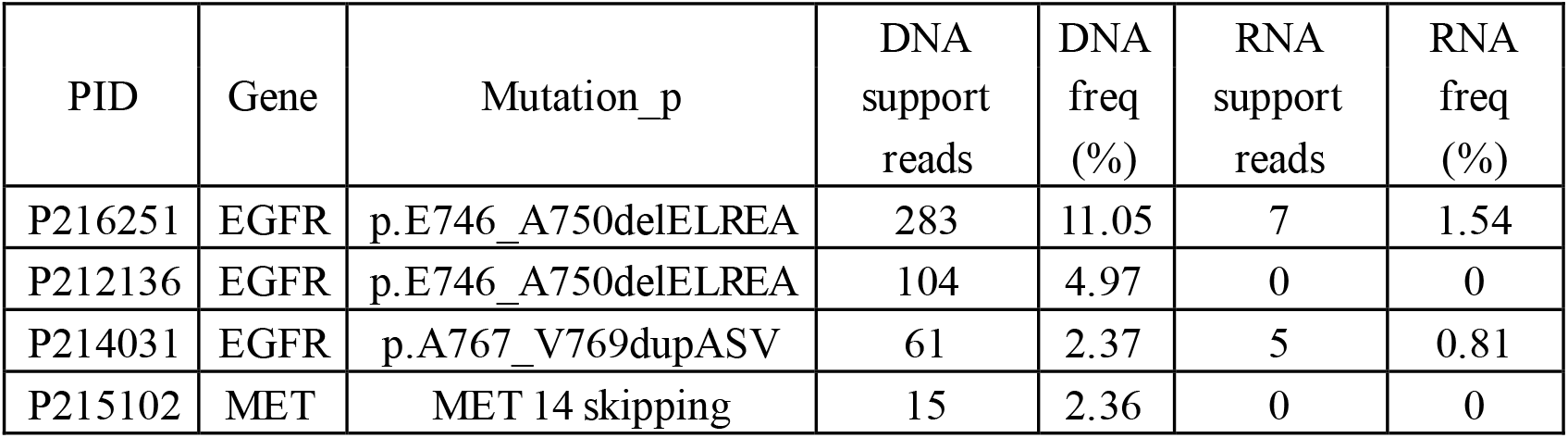
Indels missed at the RNA level

| PID | Gene | Mutation_p | DNA support reads | DNA freq (%) | RNA support reads | RNA freq (%) |
| --- | --- | --- | --- | --- | --- | --- |
| P216251 | EGFR | p.E746_A750delELREA | 283 | 11.05 | 7 | 1.54 |
| P212136 | EGFR | p.E746_A750delELREA | 104 | 4.97 | 0 | 0 |
| P214031 | EGFR | p.A767_V769dupASV | 61 | 2.37 | 5 | 0.81 |
| P215102 | MET | MET 14 skipping | 15 | 2.36 | 0 | 0 |

**Table 11.** Indels detected only in RNA

| Sample ID | Gene | Mutation_p | DNA support reads | DNA freq (%) | RNA support reads | RNA freq (%) |
| --- | --- | --- | --- | --- | --- | --- |
| P219198 | EGFR | p.L747_A750delinsP | 0 | 0 | 40 | 2.42 |

#### Measuring gene expression with the RNA panel

The standard material Tumor V2 was diluted 20X, 50X, 75X, 100X, and 125X with GM12878 FFPE RNA. For the same dilution, 50 ng, 100 ng, and 200 ng were used for library construction and RNA panel capture, which were repeated three times. The coefficient of variation (CV) value of gene expression for different starting amounts of RNA at the same dilution was calculated, and the results showed that the median value of the CV <=30% (Fig. 4). This proves that the RNA panel has good repeatability.

**Fig. 4.**
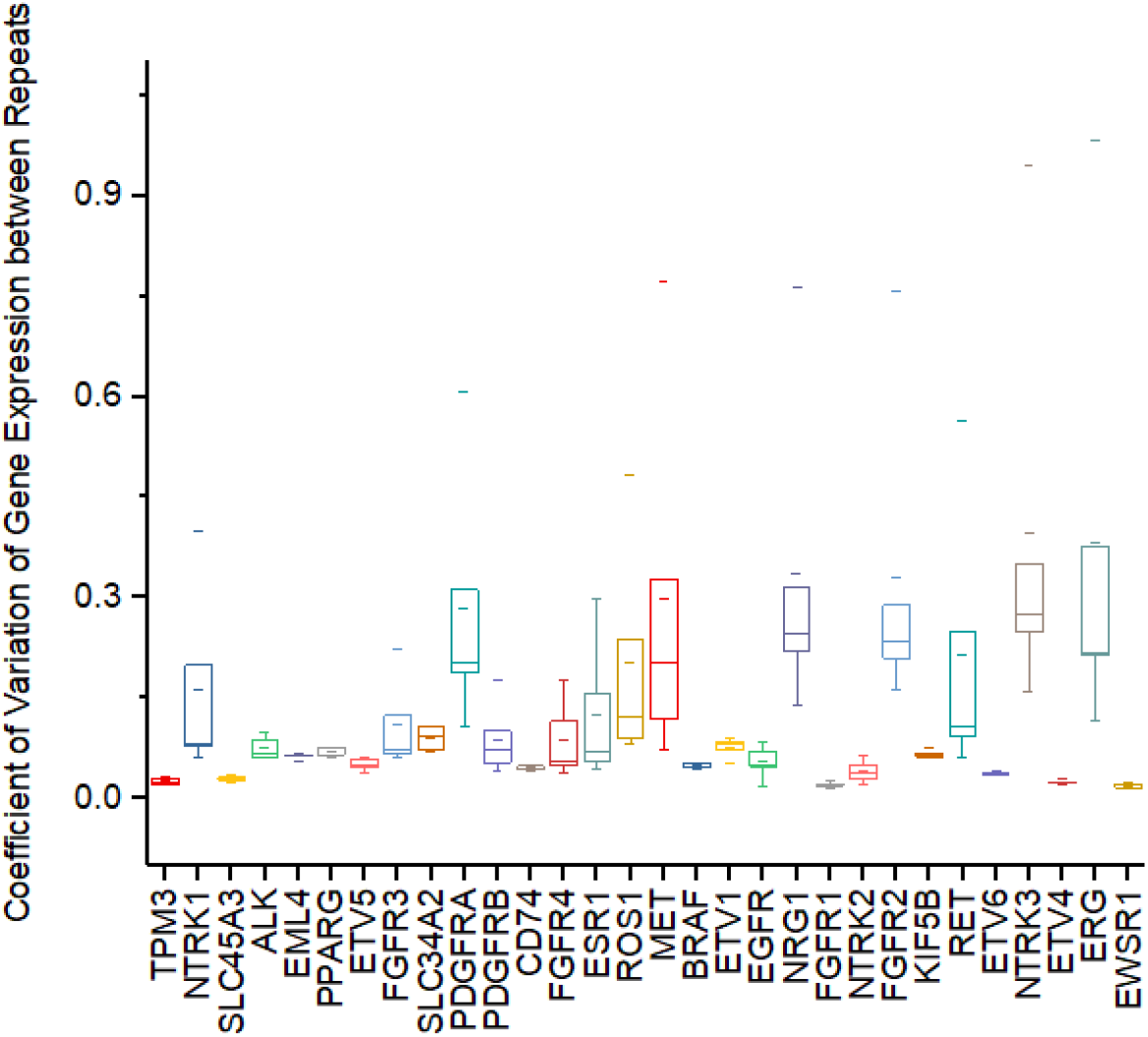
The CV value of gene expression for different starting amounts of RNA at the same dilution

The RNA panel experiment was performed on 16 clinical FFPE samples and repeated twice. The correlation coefficient of the expression of genes with target drug-related changes between the two replicates was greater than 0.9. This proves that the RNA panel has good repeatability for clinical sample gene expression.

Thirteen clinical FFPE samples were subjected to RNA-seq and RNA panel experiments to analyze the consistency of the expression levels of target drug-related genes on the two technology platforms. The correlation coefficient for the expression of 15 genes (EGFR, ESR1, FGFR1, FGFR2, FGFR3, FGFR4, MET, NRG1, NTRK1, NTRK2, NTRK3, PDGFRA, PDGFRB, RET, ROS1) between the RNA panel and RNA-seq was greater than 0.9 (Fig. 5).

**Fig. 5.**
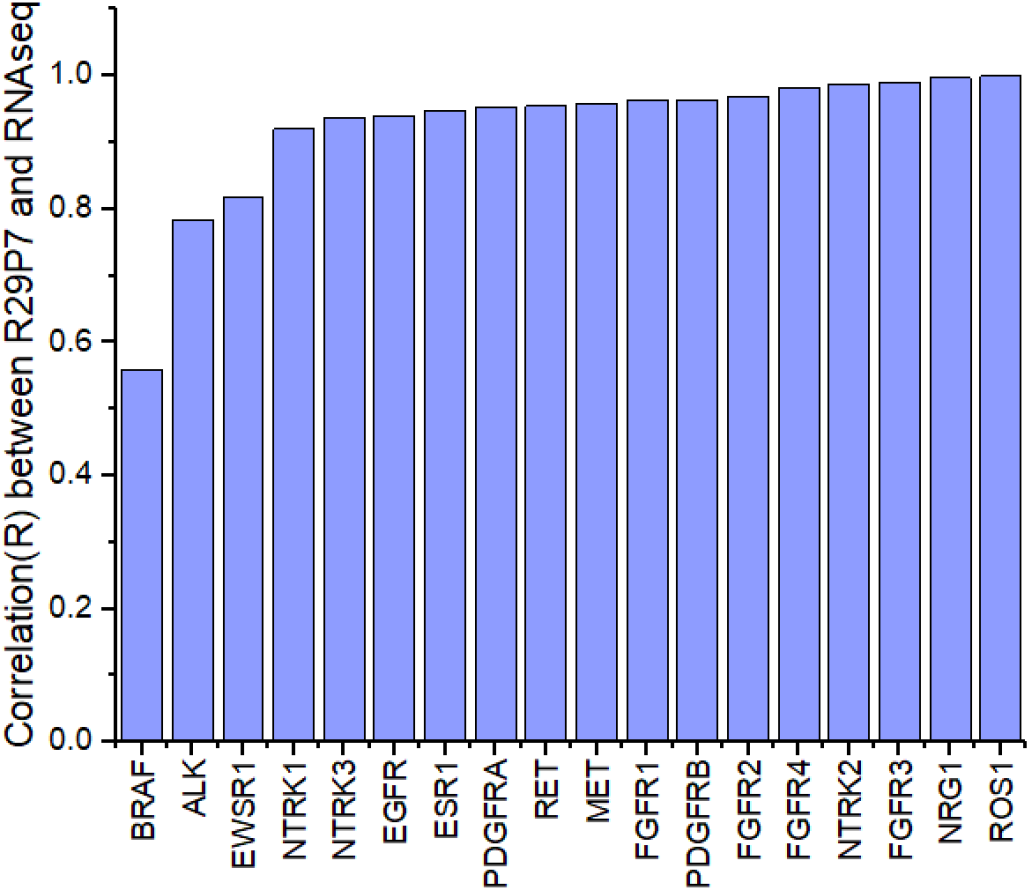
Correlation of gene expression detected by RNA panel and RNA-seq

#### Correlation of gene copy number (GCN) and RNA reads per kilobase per million mapped reads (RPKM)

The DNA copy number and RNA expression of the EGFR gene in 1253 FFPE samples was positively correlated (R=0.8) (Fig. 6).

**Fig. 6.**
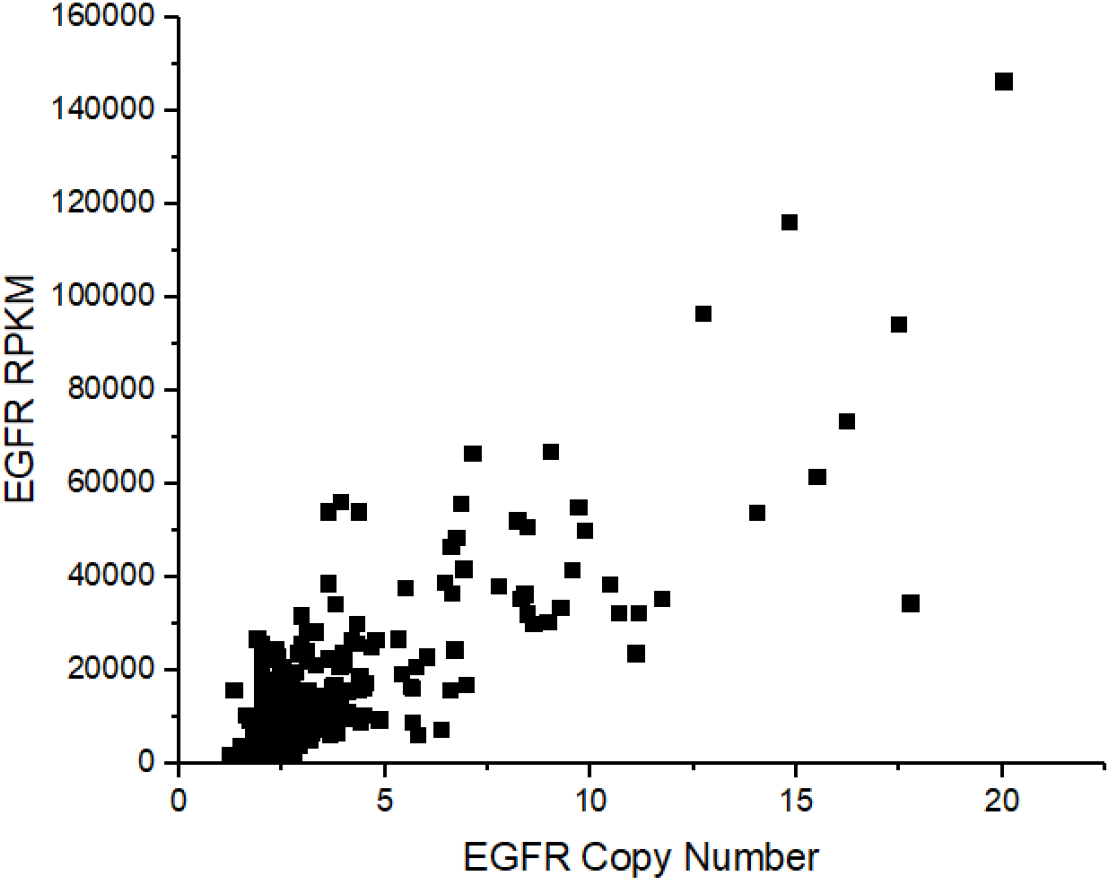
Correlation of EGFR copy number and RNA RPKM

When assessing the MET DNA copy number and MET gene expression RPKM value of 1253 samples (Fig. 7), it was found that a high DNA copy number (GCN≥6) corresponded to high gene expression, but the DNA copy number of samples with high gene expression was not always high.

**Fig. 7.**
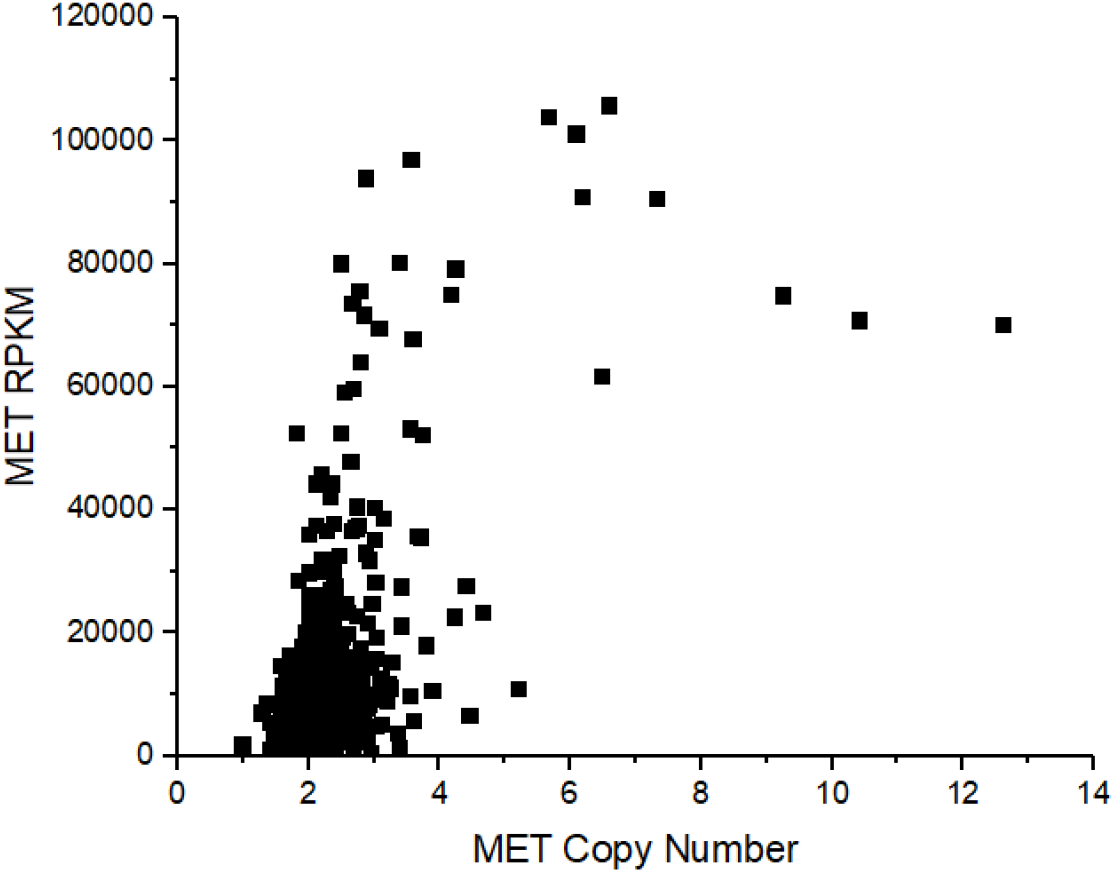
Correlation of MET copy number and RNA RPKM

Twenty-one FFPE samples were used for the detection of MET IHC and RNA expression levels. The MET gene expression level of IHC 3+ samples was higher than that of IHC 1+/2+ samples. The statistical test showed that the gene expression levels of the two groups were significantly different (Fig. 8). Twenty-seven FFPE samples were tested for EGFR IHC and RNA expression. The MET gene expression of IHC 3+ samples was higher than that of IHC 1+/2+ samples, but the difference in gene expression between the two groups was not significant (Fig. 9). The number of IHC samples in this study was small, so there was no definite conclusion, but it is possible to use gene expression to supplement the existing gene amplification methods FISH and IHC.

**Fig. 8.**
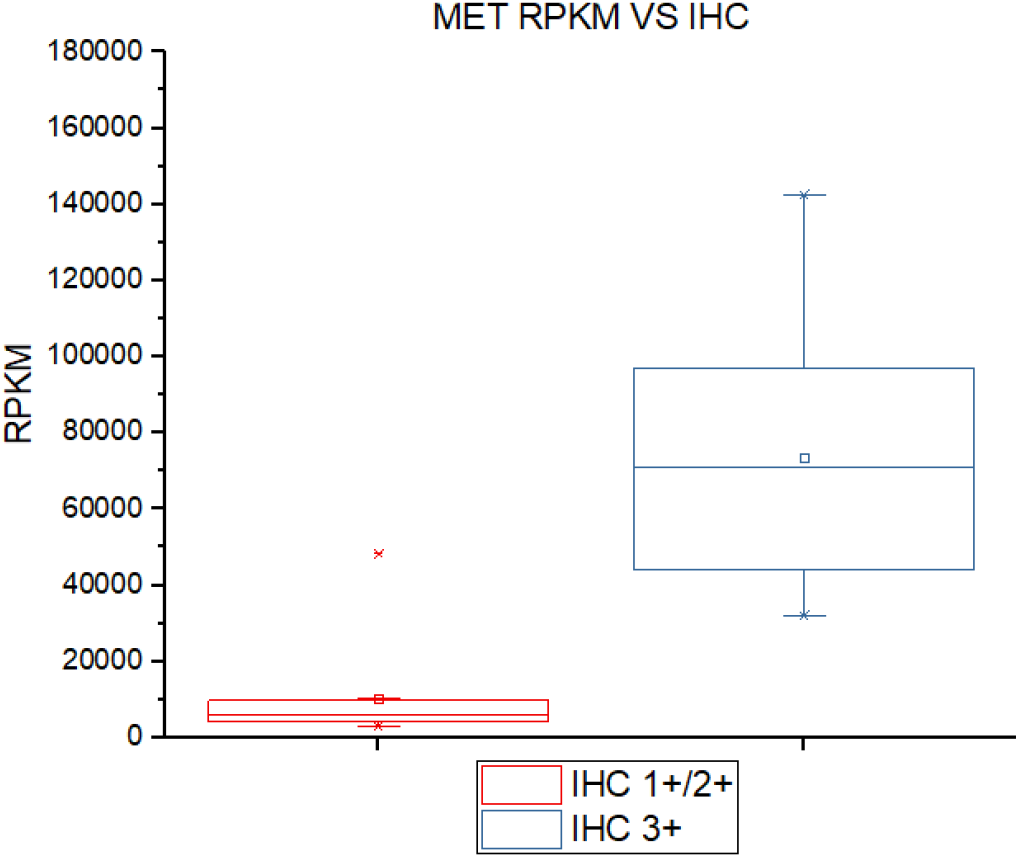
Correlation of MET IHC staining and gene expression

**Fig. 9.**
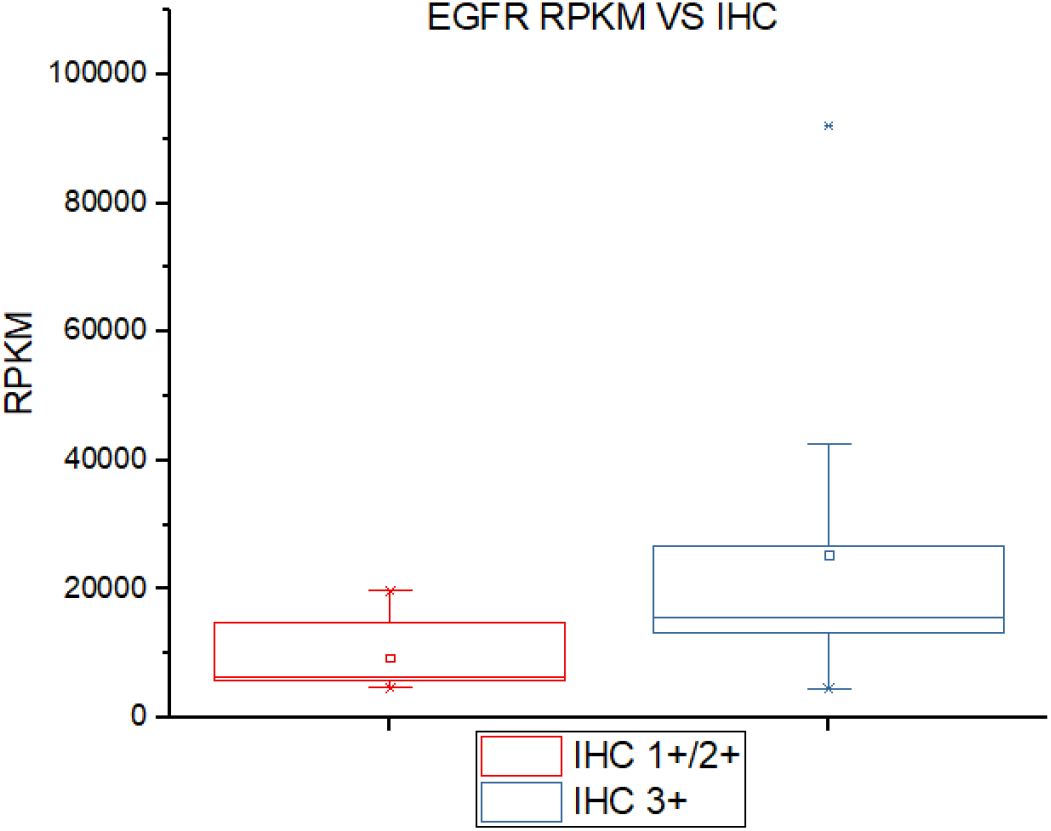
Correlation of EGFR IHC staining and gene expression

## DISCUSSION

Compared to previous methods to identify gene fusion, such as fluorescence in situ hybridization (FISH) and immunohistochemistry (IHC), next-generation sequencing (NGS) has greater power in massively parallel sequencing as well as the ability to determine specific breakpoints and identify partner genes. Testing based on the DNA panel is designed to cover both exons and introns of the target genes so that the complexity of gene fusion and rearrangements can increase the difficulties of testing, especially when intergenic regions are involved. In addition, the results of sequencing based on the DNA panel can affect the detection of gene fusions and the evaluation of potential therapies due to the complex processes of transcription and posttranscriptional regulation. In contrast, the RNA panel has unique advantages because it only detects regions of transcribed exons, excluding introns and intergenic regions. Although reports have shown that complex rearrangement with ALK, ROS1 and RET can be detected by DNA-based NGS and most of the detected gene fusions are sensitive to targeted drugs ^10–12^, the results from DNA-based NGS fail to provide direct evidence to show whether those rare fusions can generate transcribed products with desired functions^13^. Therefore, independent tests, such as IHC, are required to help determine drug sensitivity. An RNA transcript is a product transcribed by a sequence of gene fusion. It is able to provide more reliable evidence for predicting the response to targeted therapy. In this study, DNA-based NGS detected seven complex rearrangement, 7 of which were verified as transcripts of gene fusion by RNA-based NGS (4 ALK fusions, 2 RET fusion, and 1 FGFR fusion).

There are more than 20 types of EML4-ALK fusion variants. The most common variants are V1 and V3a/b (60%). Better responses to crizotinib were reported in non-small cell lung cancer (NSCLC) patients with the V1 variant than in those with non-V1 variants^14–16^. The results of this study showed that the relative RNA expression of EML4-ALK fusion V1 was significantly higher than that of EML4-ALK fusion V3 detected by RNA NGS. This may be one of the reasons why NSCLC patients with EML4-ALK fusion V1 have a better response to crizotinib than those with V3. The level of fusion gene RNA expression may be a maker for predicting the effect of medication in patients.

The results for METex14 skipping mutation at the DNA level are different from those at the RNA level. In the study, 26 cases of METex14 skipping were detected at the RNA level, whereas only 21 cases were detected at the DNA level; no mutation was detected in the DNA splicing region in the other 5 cases. A possible reason is the low frequency of the splicing mutation. Studies have shown that mutations can occur in the splice acceptor site, branching site and polypyrimidine tracks. It is a technical challenge to design panels that cover those regions. Another possible reason is that the METex14 skipping mutation was caused by the deletion of long DNA fragments at the DNA level^17^, such as fragments covering the entire exon 14, so that the designed panel failed to detect these noncanonical mutations. At the RNA level, probes are designed to cover the MET transcript only. Because the METex14 skipping mutation causes deletion of the entire exon 14, RNA-based testing can indicate the presence of METex14 skipping if exon 13 is detected to be followed by exon 15 instead of exon 14. The panel covers fewer sequencing regions, so it is unlikely to miss any mutation.

In this study, several cases of rare fusions involving ALK, RET and ROS1, such as CLTC-ALK, KLC1-ALK, LMO7-ALK, DCTN1-ALK, KIF5B-ALK and PICALM-RET, were detected by RNA NGS. By analyzing the results from the testing of rare gene fusion, we noticed that DNA testing is limited if the targeted regions contain long introns, repeated sequences, or genes with special structures (MET/ROS1/NTRK). It is also unfavorable for detecting complex fusions or rearrangements. Additionally, the final transcribed products may vary due to the complex mechanism of transcription and the stochasticity of alternative splicing, therefore affecting the evaluation of the benefits of potential targeted therapy. In this case, RNA-based NGS can increase the rate of fusion detection and provides more treatment opportunities for patients with more accurate information.

Previously, RNA was only used for the detection of gene fusions because the results of RNA sequencing (RNA-seq) were considered to be inaccurate for other types of gene mutations. However, most research related to the detection of RNA single-nucleotide variations (SNVs) uses whole-transcriptome sequencing to explore the detection of RNA SNVs. Comparing whole-exome sequencing (WES) to whole-transcriptome sequencing in terms of the detection of SNVs in BRAF and RAS genes in tumor tissues, the study showed that 94% of the SNVs (variant allele frequency (VAF) >10%) detected by WES were successfully detected by RNA-seq, but only 11% of SNVs (VAF = 5-10%) could be detected by RNA-seq. The results demonstrated that whole-transcriptome analysis with total RNA-seq had a low sensitivity to SNV mutations^6^. In this study, we collected over 1000 clinical NSCLC tissue samples and compared the results from targeted DNA and RNA NGS in parallel. The testing results from the RNA NGS were highly consistent with those from the DNA NGS (PPA >97%). Therefore, targeted RNA-seq was applicable and sensitive for detecting SNVs/insertions/deletions (indels) that were related to targeted drugs. The high sensitivity of RNA-seq could probably be attributed to the RNA panel we used for capturing the targeted RNA in the research. In the RNA panel, the probes were designed to enrich critical genes that were related to targeted drugs so that the sequencing depth was much deeper than that of the whole-transcriptome sequencing (Fig. 10). Therefore, the sensitivity of RNA panel sequencing is higher than the sensitivity of whole-transcriptome sequencing in terms of the detection of SNVs.

**Fig. 10.**
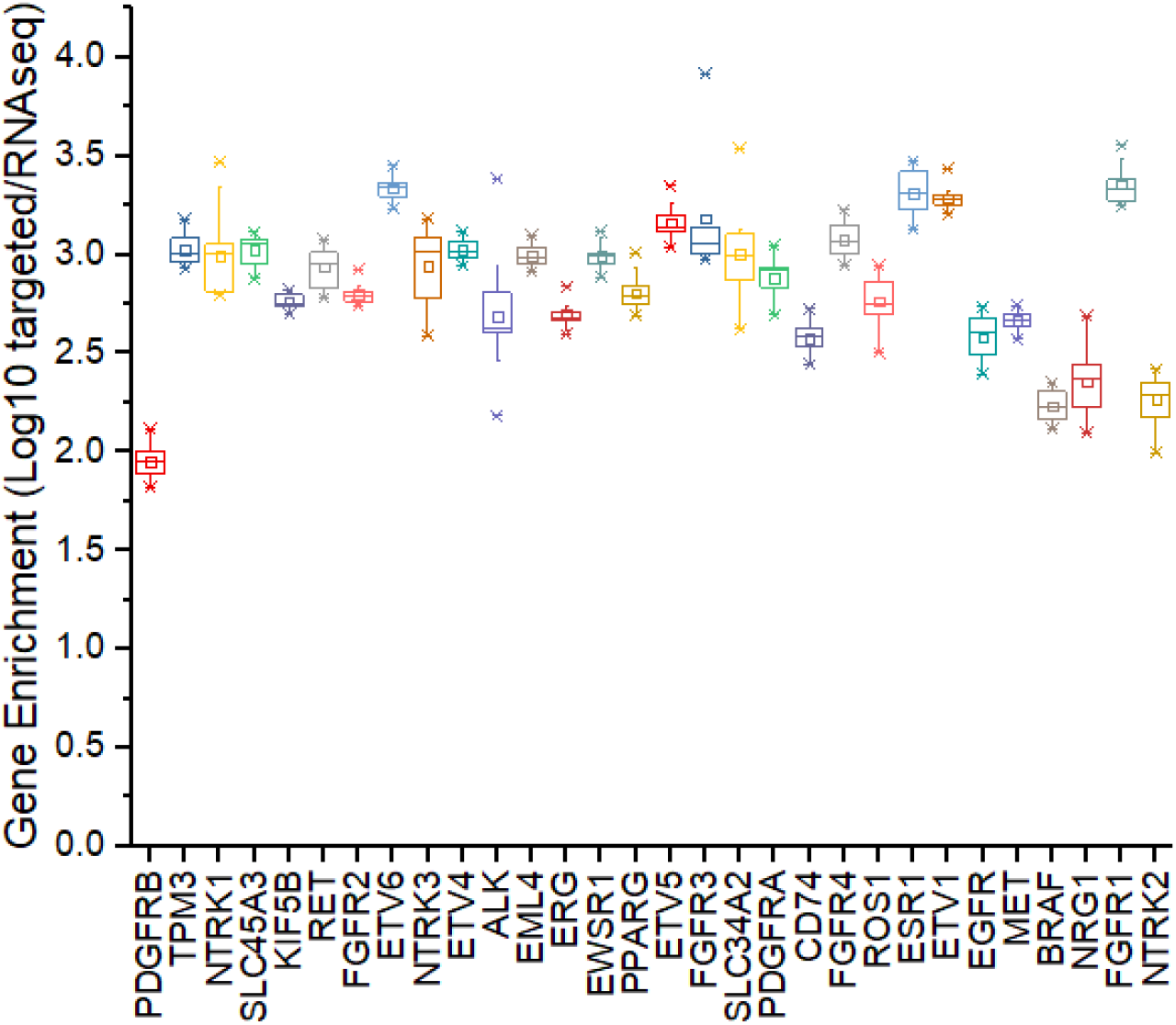
Boxplot of targeted RNA panel enrichment for genes included in the panel

FISH and IHC are the two commonly used assays to detect copy number variations (CNVs). As NGS technology has rapidly developed, DNA NGS has also been applied for CNV detection.

However, there is no consensus on the criteria for determining the MET gene amplification on subsequent medications in NSCLC. Studies have shown that after the uptake of capmatinib in NSCLC patients, patients with MET gene copy number (GCN) ≥ 6 benefited from the medication the most, with a median progression-free survival of 9.3 months. In comparison, the progression-free survival (PFS) of patients with IHC MET+++ was 7.3 months; the PFS of the patients with MET/CEP7 ratio ≥2.0 was 5.6 months; the PFS of the patients with an H-score between 150 and 300 was 3 months; and the PFS of the patients with an H-score ≥≥300 was 7.3 months^18^.

We collected over 1000 NSCLC cases to study the DNA copy numbers and the level of gene expression. We found that the correlations between the DNA level and the RNA level were different for the EGFR and MET genes. For the EGFR gene, the copy number at the DNA level was positively correlated with the copy number at the RNA level (R=0.8). The results were consistent with the results from previous studies. Researchers studied the relationship between EGFR copy number and protein expression in NSCLC by IHC and FISH and found that protein expression levels were significantly correlated with the GCN per tumor cell (p <0.001)^19–20^. In contrast, for the MET gene, the level of RNA expression was high when the level of DNA copy number was high (GCN≥6). However, the RNA level was also high in some cases when the DNA level was low. This was also consistent with the results from previous studies. Researchers who studied the relationship between MET GCN and IHC H-score in NSCLC found that the H-score was greater than 100 in the samples with GCN > 6, but in many samples with GCN <6, the H-score was also greater than 100^21^. We speculated that MET gene amplification is not the only factor that leads to an increase in MET expression. Other causes could be the regulation of transcription^22^, fusion mutations, etc.

