## supplemental for "Detection of Multiple Types of Cancer Driver Mutations Using Targeted RNA Sequencing in NSCLC": Supplemental Figures.docx

a．ALK-ERC2 (A19; E8)

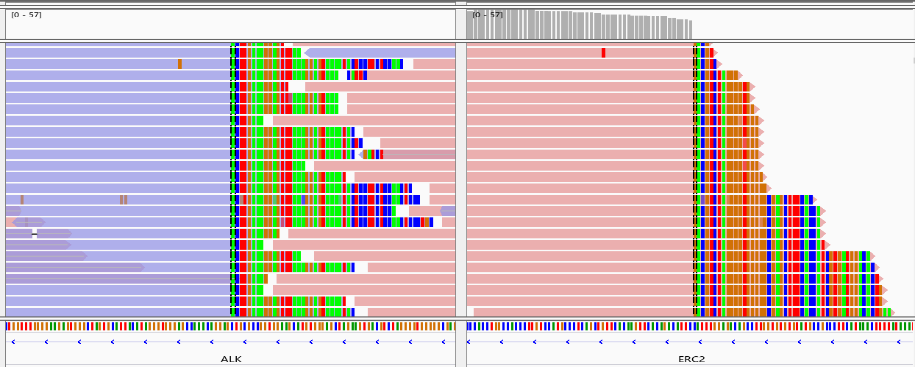

b. EML4-ALK V1

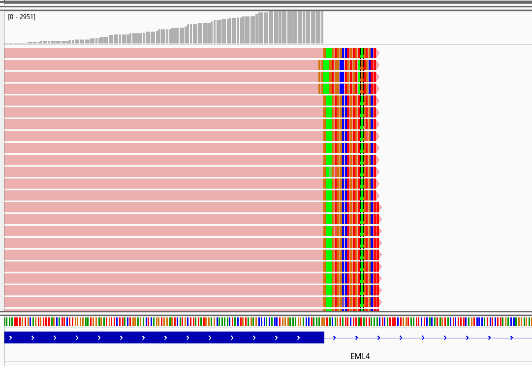

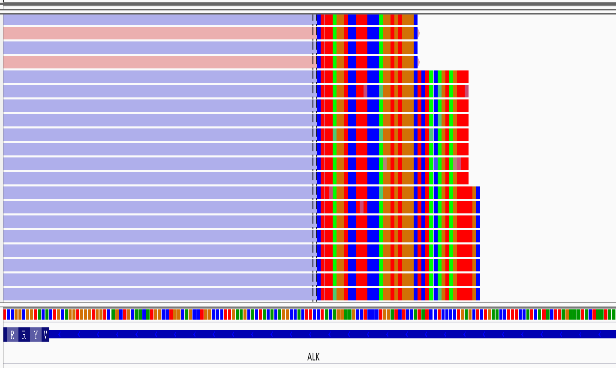

c.

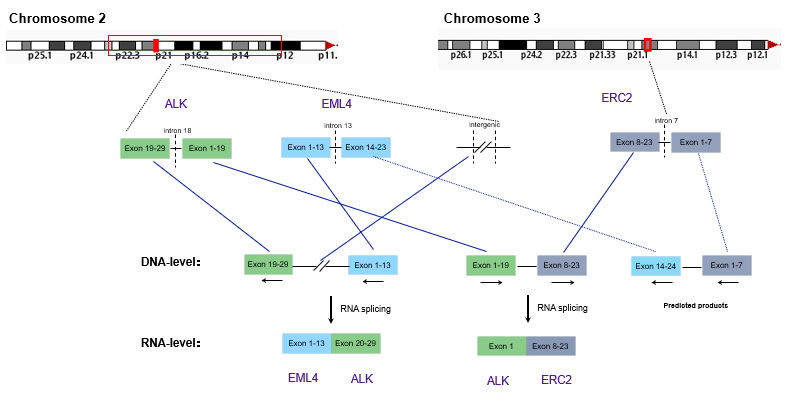

**Fig. S1.** A complex structural rearrangement. Picture a shows ALK-ERC2 (A19; E8) fusion at the DNA level. Picture b shows the EML4-ALK V1 fusion at the RNA level. Picture c shows a possible process of gene rearrangement.
