## supplemental for "Detection of Multiple Types of Cancer Driver Mutations Using Targeted RNA Sequencing in NSCLC": Supplementary Material.docx

**Generation of the DNA panel**

The DNA panel covers the genes related to 25 common solid tumors. It is capable of detecting 790 genes located in the exome and other hotpot regions in tumor tissues through the NGS platform. The target region is 2189 kb long and covers loci related to targeted therapy, immunotherapy and chemotherapy. The capture probes were customized by Integrated DNA Technologies (IDT) for the detection of gene mutations, including gene fusion, single-nucleotide variations (SNVs) and copy number variations (CNVs).

Extracted from tissue samples, DNA was captured and sequenced using the DNA panel. Data analysis showed that the map ratio was 0.999±0.002, the target ratio was 0.767±0.026, and the coverage 100x was 0.999±0.001. All the samples generated enough data for subsequent analysis.

**Validation of SNV mutations with the DNA panel**

To evaluate the sensitivity, specificity and positive predictive value (PPV) of the DNA panel in the detection of SNVs, DNA from a variety of clinical formalin-fixed, paraffin-embedded (FFPE) samples was mixed to prepare 20 FFPE standard pools with distinct frequencies of mutations. The DNA panel was used to capture the standard library. The results from the analysis of SNV detection showed the sensitivity, specificity and PPV performance of the DNA panel (Table S1).

Table S1. Summary of analytical performance of SNV measured by DNA panel

| Type | De-dup  depth | MAF | Sensitivity  (95% CI) | Specificity  (95% CI) | PPV  (95% CI) |
| --- | --- | --- | --- | --- | --- |
| Hotspot | >500 | >2% | 96.77%  (88.97%-99.11%) | 100.00%  (99.99%-100.00%) | 100.00%  (96.23%-99.88%) |

To evaluate the accuracy of the DNA panel for SNV detection, we used 40 FFPE samples from external quality assessment (EQA) and 4 clinical tumor tissue samples with confirmed mutations by droplet digital PCR (ddPCR). The DNA was extracted from the samples and used to construct a library using the same operation procedures as that of the tissue samples for the DNA panel. After capture and analysis, the NGS results were compared to the set threshold, and the sensitivity, specificity and PPV were all 100%.

Table S2. Summary of clinical performance of SNV measured by DNA panel

| Sensitivity  (95% CI) | Specificity  (95% CI) | PPV  (95% CI) |
| --- | --- | --- |
| 100.00%  (90.36%-100.00%) | 100.00%  (87.54%-100.00%) | 100.00%  (90.36%-100.00%) |

To evaluate the repeatability of the DNA panel for SNV detection, we used 4 gDNA samples from clinical tumor tissues and repeated the experiment 3 times. The repeatability of 4 loci of interest in the 3 repeats was CV ≤ 20%, indicating the stability of the DNA panel for detection.

**Validation of fusion mutations with the DNA panel**

To validate the sensitivity, specificity and PPV of the DNA panel for fusion detection, 8 cell-line reference samples, including 7 positive control samples and 1 negative (12878) control sample, were used, and the mutations were determined by ddPCR. The fusions found in the controls were SLC34A2-ROS1 (E4-E32), EML4-ALK (E6-E20) and EML4-ALK (E13-E20), with frequencies between 0.00% and 3.4% (Table S3). The standard product was taken with an initial amount of 50 ng. We analyzed the results of fusion detection with the DNA panel, and the performance of the DNA panel in fusion detection was evaluated in terms of the sensitivity, specificity and PPV.

Table S3. Summary of analytical performance of fusion measured by DNA panel

| Expectation | Sensitivity  (95% CI) | Specificity  (95% CI) | PPV  (95% CI) |
| --- | --- | --- | --- |
| >0.001 | 89.86%  (80.51%-95.00%) | 100.00%  (84.54%-100.00%) | 100.00%  (94.17%-100.00%) |

**Validation of CNVs with the DNA panel**

To evaluate the sensitivity, specificity and PPV of CNV detection with the DNA panel, 8 cell-line reference samples, including 7 positive control samples and 1 negative (12878) control sample, were used, and the mutations were determined by ddPCR. The genes involved in the CNVs in the controls were ERBB2 and MET (2<CN<4) (Table S4). The targets in the standard product were captured by the DNA panel, and the testing results of the CNVs were analyzed. The performances of the DNA panel in CNV detection was assessed according to the sensitivity, specificity and PPV.

Table S4. Summary of analytical performance of CNV measured by DNA panel

| CNV | Sensitivity  (95% CI) | Specificity  (95% CI) | PPV  (95% CI) |
| --- | --- | --- | --- |
| >2.2 | 100.00%  (97.04%-100.00%) | 100.00%  (93.36%-100.00%) | 100.00%  (97.04%-100.00%) |

To evaluate the repeatability of CNV detection with the DNA panel, we used 4 clinical tumor tissue samples with different tumor percentages and repeated the experiment 3 times. The repeatability of copy number detection for the ERBB2 and MET genes was evaluated (CV ≤ 2%), and the overall consistency was 100% compared to the set threshold.
